# Failure to meet the exchangeability assumption in Bayesian multispecies occupancy models: Implications for study design

**DOI:** 10.1101/2025.04.30.651473

**Authors:** Gavin G. Cotterill, Douglas A. Keinath, Tabitha A. Graves

## Abstract

Bayesian hierarchical models are ubiquitous in ecology. Random effect model structures are often employed that treat individual effects as deviations from larger population-level effects. In this way individuals are assumed to be ‘exchangeable’ samples. Ecologists may address this exchangeability assumption intuitively, but might in certain modeling contexts ignore it altogether, including in situations where it may have large implications for study design. Multispecies occupancy models based on detection/non-detection data are an approach that can be utilized by those tasked with monitoring rare and endangered species because most literature suggests that, compared to single species occupancy models, improved parameter estimates are assured. Yet, we illustrate through a power analysis how sampling requirements to detect experimental treatment effects vary tremendously depending on whether the species exchangeability assumption is met. The degree to which species in a community respond similarly to covariates governs the ability to accurately estimate parameters using multispecies occupancy models. Detecting small or moderate changes in occupancy resulting from habitat restoration treatments may be impossible for small datasets (e.g., < 36 sampling locations, each surveyed < 8 times) even with a paired treatment-control design if the exchangeability assumption is violated. By contrast, when the assumption is met, small effects may be confidently estimated with as few as 12 sampling locations (6 pairs) and 6-8 survey events. Often, it may be impossible to know whether the exchangeability assumption is met. The statistical power needed to accurately estimate species-specific effects using detection/non-detection multispecies occupancy models depends on the unknown values of treatment effects and whether responses by species in the community diverge. When the species exchangeability assumption is violated, and at lower levels of sampling effort, multispecies occupancy models may provide worse inference than single species occupancy models.

## Introduction

Identifying group relationships, as in individuals within a population or functional groups within a species community, is a central challenge in statistical inference and ecology (Draper et al., 1993; Elton, 1946; Hubbell, 2005). ‘Exchangeability’ can be broadly thought of as the ability to make inference on new individuals based on a sample of a population (Lindley & Novick, 1981). Absent additional information, model assumptions like exchangeability are defensible (Gelman et al., 2013) but can have unforeseen consequences. A classic example of how failure to understand group membership results in improper inference (due to one or more confounding variables) is Simpson’s paradox (Simpson, 1951). For example, consider a model where the dependent variable is a patient’s recovery time. The most obvious independent variable may be the disease with which patients are afflicted, but other important groupings could include demographic information such as age, sex, underlying medical conditions or socioeconomic status – omitting any of these might drastically alter inference and prediction error when applied to new patients (Greenland & Robins, 1986). In modeling terms, within-group variation can be accommodated through a mix of random intercept and random slope terms (Muff et al., 2020). Similarly, among-group variation can be accommodated through changes to the hierarchical model structure. In both cases, the value of added model complexity, assuming there is sufficient data to fit the model, still rests upon an assumption that individual-group relationships are justified.

For many ecological questions, grouping justifications seem obvious. For instance, the natural histories of ungulates are well-documented. If investigating resource selection of a single species like mule deer (*Odocoileus hemionus*), a reasonable strategy would be to employ a random effect structure that nests individuals within regional subpopulations while also accounting for seasonal variation (winter vs summer habitat) and sex (especially during fawning season). In this way, individual selection patterns are treated as deviations drawn from a group-level distribution based on sex, season, and region. Doing so in a Bayesian context improves parameter estimates while taking a fuller account of parameter uncertainty (Ellison, 1996). In contrast, modeling the resource selection of multiple ungulate species together (e.g., mule deer, moose [*Alces alces*], bighorn sheep [*Ovis canadensis*]) might be met with skepticism, even after taking species into account, based on their divergent habitat preferences and life histories. Yet, there are modeling frameworks where similar choices are the norm and exchangeability is not always explicitly addressed.

Ecotoxicology risk assessments for chemical substances provide an example where species exchangeability has often been violated. These models historically assumed, often incorrectly, that tolerances of all species in a community were exchangeable for each new substance (Craig et al. 2012). The issue has been addressed with various model extensions and diagnostic tests but still depends heavily on domain-specific knowledge and large, published datasets that characterize species-specific responses to substances (Migliorati et al., 2021). Multispecies occupancy models (MSOMs) make a very similar assumption – that all species in a community are exchangeable for all environmental or detection covariates (i.e., the species exchangeability assumption [SEA]; Kéry and Royle 2015).

Occupancy models simultaneously estimate the probability that a species occurs at a specific location and the probability that the species is detected (MacKenzie et al., 2003). The models can operate on data from a single species (single species occupancy models; SSOMs) or on data from multiple species collected during the same sampling event (community, or multispecies, occupancy models; Dorazio and Royle 2005). SSOMs make fewer assumptions than MSOMs and are therefore more broadly applicable. However, precise parameter estimates using SSOMs can require many sampling events across multiple locations. Acquiring sufficient data for rare or cryptic species can be especially challenging, and though collecting data on multiple species can create additional work, in practice, it does not always increase project costs (e.g., bird point counts, eDNA samples of water, air, or flowers, vegetation transects). MSOMs can therefore be appealing because the precision and accuracy of parameter estimates for hard-to-detect species are improved by sharing information with more frequently observed species (Zipkin et al., 2009), thus reducing the overall level of required sampling (i.e., partial pooling, also referred to as “borrowing strength”; Gelman and Hill 2007). MSOMs accomplish this through their random effect structure (community-level average effects inform species-level effects) but this data sharing is only appropriate and useful if species are exchangeable (Pacifici et al., 2014).

Several solutions have been explored to address the existence of multiple groups in community datasets. One approach adds levels to the model hierarchy *a priori* that correspond to species-relatedness (Ives & Helmus, 2011). This may be appropriate in some, but not all, contexts. Closely related species are often morphologically, physiologically, and behaviorally similar, and competition theory suggests that the persistent co-occurrence of similar species requires resource partitioning (Chesson, 2000; Hardin, 1960). This can include spatial partitioning resulting from competitive exclusion (Durant, 1998; Sollmann et al., 2012) which would violate the SEA of MSOMs for occupancy probabilities. Likewise, temporal partitioning, where similar species may be more or less detectable during certain periods of the diel cycle (Hayward & Slotow, 2009; Stewart et al., 2014) or season (Scriven et al., 2016), could violate the exchangeability assumption for detection probabilities depending on the sampling method. An alternate solution derives functional groupings based on multiple species-specific traits (Ovaskainen et al., 2017) but requires additional prior knowledge – information that may be unavailable for many study systems. More recently, latent-class model extensions to MSOMs were proposed to directly estimate functional groups based on similarities of species’ responses to covariates (Sollmann et al., 2021), but these models suffer from additional identifiability issues and are not well-suited to sparse, binary data (Qiu & Yuan, 2024). How species respond to environmental covariates is often precisely the question of interest – and yet also the information that should ideally guide grouping decisions. This is a particularly vexing issue when designing studies, i.e., determining the amount of survey effort needed to guide adaptive management of understudied species or communities.

As a motivating example, we consider a hypothetical pollinator community in the western United States where native pollinators are declining due to complex causes including loss of floral resources (Ogilvie et al., 2017), increasing temperature and drought, and neonicotinoid pesticide use (Janousek et al., 2023). Although directly addressing some of these factors may fall outside land managers’ purview, improving habitat quality can help mitigate stressors. Some experimental treatments currently implemented on public lands in the region are expected to benefit all species in the pollinator community – e.g., restoration plantings to increase native flowers (Drobney et al., 2020) – while other treatments are expected to benefit some species at the expense of others – like rotational grazing of livestock that enhances habitat for ground-nesting bees (Goosey et al., 2024). Despite the ubiquity of habitat restoration and conservation actions, rigorous assessment of outcomes is lacking (Binley et al., 2025; Sutherland et al., 2004). In our own research, we found little guidance for practitioners interested in quantifying treatment effects on species of concern or broader communities with detection/non-detection data under the resource constraints common to public land management agencies (Moore et al., 2011; Smiley, 2008). We therefore performed a simulation study to identify minimum data requirements to accurately quantify species-specific treatment effects using detection/non-detection data. We hypothesized that actions producing ‘winners and losers’ would violate the species exchangeability assumption of MSOMs and require much larger datasets to accurately estimate species’ responses. We compare MSOM results against more conservative SSOMs using several error metrics. Our goal was to inform sample designs to evaluate restoration actions that could be applied regardless of the taxa of interest or data collection method.

## Materials and methods

We analyzed known species richness models which consider a predetermined number of community species. This choice was made for several practical reasons: (1) Our focus was not on estimating species richness or biodiversity metrics; and (2) Pollinator community sampling – whether human surveyors, bowl traps, eDNA – often includes visually or genetically similar species leading to a mix of species-, genus-, or family-level identifications due to bottlenecks in taxonomic expertise, methodological precision, and cost (Woodard et al., 2020) that restrict the depth of taxonomic identifications. Understanding how various species or functional groups respond might be more important than biodiversity metrics when management objectives include monitoring species of concern.

### Model formulations

The multispecies occupancy model (MSOM; Dorazio and Royle 2005) treats the latent occupancy (i.e., unobserved ecological truth or state process; *z*) at site *i*, for species *k*, as a Bernoulli random variable based on the probability *ψ*_*ik*_,

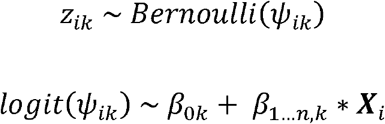

where *ψ*_*ik*_ is logit-transformed to accommodate a linear model estimating a random effect of species identity (the intercepts *β*_0*k*_) and one or more environmental effects (*β*_1_…_*n*,*k*_) from a vector of site-specific covariates (***X***_*i*_). This process model is linked to an observation model that estimates species’ detection probabilities (*p*_*k*_). In it, data (*y*_*ijk*_; i.e., whether species *k* was observed at site *i* during sampling event *j*) is a Bernoulli random variable based on probability *θ*_*ijk*_, which is the probability of observing the species conditional on its occupying the site and its probability of detection:

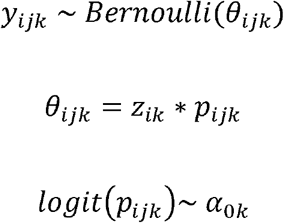

Detection probabilities can also vary as a function of site or survey event (implied by the inclusion of three subscripts in the equation above), but here we simulated from, and subsequently fit, an intercept-only (*α*_0*k*_) model in which detection probabilities only varied by species. This reflects cases in which surveys occur under ideal conditions, a common practice for pollinator surveys (FWS, 2019), reduces the dimensionality of the power analysis, and speaks to the question of minimum data collection needs (if surveying during conditions that reduce detection rates, more data will be required). In our process model we considered a single habitat covariate: that of categorical treatment effect. In this case, ***X***_*i*_ represents a ‘dummy variable’, the vector of zeros or ones indicating whether a site belonged to the control (0) or treatment (1) group. On their own, the *α*_0*k*_, *β*_0*k*_ and*β*_1*k*_ parameters are of some interest, but our main goal was to estimate the change in species-specific occupancy probabilities resulting from a treatment effect 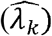, which were the derived quantities,

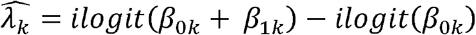

where *ilogit* is the inverse logit function 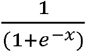.

This hierarchical model uses priors to control the species-level random effects as required by Bayesian models (Kéry & Royle, 2015). In ecological research, priors are usually uninformative or weakly informative distributions (Northrup & Gerber, 2018). For example, priors for *α*_0*k*_, *β*_0*k*_ and *β*_1*k*_ could all be normal distributions with a zero mean and some error. Doing so is equivalent to fitting many single-species occupancy models all at once. The difference in the MSOM is that priors for *α*_0*k*_, *β*_0*k*_ and *β*_1*k*_ are drawn from community-level distributions for each of the parameters (called hyperpriors). The prior structure for a single parameter therefore took the form:

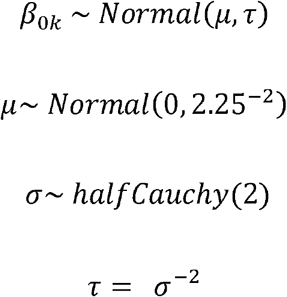

Here species-specific occupancy intercepts (*β*_0*k*_) were drawn from normal distributions governed by the community-level mean (*μ*) and precision (*τ*). In turn, the prior for *μ* was an uninformative normal distribution, and *τ* was equal to 1/*σ*^2^, where the prior for *σ* was an uninformative half-Cauchy distribution. For analyses detailing what constitutes uninformative or weakly informative priors on the logit scale, and in particular for occupancy models, refer to Gelman et al. (2008), Northrup and Gerber (2018). Our full set of priors are described in the supplement. In addition to the MSOM, we also fit SSOMs and a hybrid model. The hybrid model pooled data for *α*_0*k*_, *β*_0*k*_ (random effects) but not *β*_1*k*_ (fixed effects) by excluding the community-level hyperprior for the beta coefficient corresponding to treatment. We fit all models in R version 4.4.1 (R Core Team, 2024) using JAGS (Plummer, 2003) and the jagsUI package (Kellner & Meredith, 2024). Code to perform our full analysis is available in the accompanying software release (Cotterill and Graves, 2025).

### Data simulation

We simulated communities that included 25 or 50 species (or taxonomic units). This level of diversity reflects some real-world pollinator communities (e.g., Kearns and Oliveras 2009), as well as resource bottlenecks (Woodard et al., 2020) that could constrain the size of communities monitored, but make the study more computationally-tractable. Reported occupancy probabilities for pollinators tend to range more widely than detection probabilities (Bergman et al., 2004; Janousek et al., 2023; MacIvor & Packer, 2016; McNeil et al., 2019; Nunes et al., 2024; van Strien et al., 2011). Therefore, species-specific occupancy intercepts were drawn from a uniform distribution between 0 and 1, whereas species-specific detection probabilities were drawn from a more conservative uniform distribution between 0.1 and 0.5. We evaluated two components of sample design: the number of sites and surveys, where surveys could represent sampling occasions (e.g., human observers) or the number of samples collected (e.g., eDNA) depending on the data collection method. We simulated 6, 12, 18, or 36 sites divided equally into treatment and control sites (e.g., 3 and 3 for 6 sites). The number of survey events was 2, 3, 4, 6, or 8, and all sites were surveyed the same number of times within each simulation.

Under each combination (species x sites x surveys), we simulated four treatment response scenarios. Each of our four treatment responses represented patterns that we might expect to see in pollinator communities when treatment influences species’ occupancy probabilities. The four treatment response scenarios are detailed here. (1) All species within the community responded similarly (hereafter ‘same response’). This scenario might represent pesticide application or the removal of highly invasive plant species that created monocultures of low value to pollinators. (2) The community consisted of two groups that responded in equal and opposite ways representing ‘winners and losers’ (hereafter ‘bimodal response’). This might be the most common situation and the clearest violation of the species exchangeability assumption of the MSOM. For example, areas that have been heavily grazed by livestock are expected to have more bare ground that may favor solitary ground nesting bees but reduce some species of butterflies (Goosey et al., 2024; Pöyry et al., 2004). (3) Species-specific responses were variable but drawn randomly from a zero-centered normal distribution (hereafter ‘random response’). The random response is a frequent textbook example, and a variant of the ‘winners and losers’ scenario where species-level responses are more variable. For each of these three treatment types, we also varied the mean magnitude of effect corresponding to small (<= |0.125|), intermediate (<= |0.25|), and large (<= |0.5|) changes in occupancy probability (refer to supplement for details). (4) We also simulated data with no response to treatment effect to investigate type 1 (false positive) error risk (hereafter ‘no response’). This could occur when treatments, intentionally or unintentionally, do not influence pollinator occupancy.

These conditions generated 400 unique community-design combinations that were each simulated 100 times resulting in 40,000 datasets. We fit three kinds of models (MSOMs, hybrid MSOMs, and SSOMs) for all species and calculated several error metrics (refer to *Error Calculations* section) to assess model performance. All models were run with three chains, with 6,000 iterations after burn-in (n = 2,000) thinned to every tenth sample (i.e., a total of 1,200 samples kept; 4,000 * 3 / 10). We stored effective sample sizes and Gelman-Rubin statistics 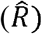 (Gelman & Rubin, 1992) for all parameters and assessed convergence by ensuring that all 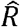 values < 1.1.

### Error calculations

We calculated error metrics for our derived quantity of interest, the species-specific estimates of treatment effect 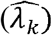, because different kinds of error and levels of uncertainty may be more important to managers in some scenarios. These included the root-mean squared error, coverage and confidence interval widths, type 1 and type 2 error risk, and what we referred to as ‘percent correctly classified’ (PCC), a simplified metric representing the degree of confidence and correct classification of the *direction* of a treatment effect. For each set of 100 replicates, to calculate root-mean squared error (RMSE) we summarized RMSE across replicates as:

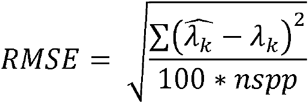

where *nspp* was the number of species in the community. Coverage was the proportion of times that λ_*k*_ was contained in the 95% or 75% credible interval for 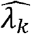 and credible interval widths were the difference between the 0.975 and 0.025 percentiles or 0.875 and 0.125 percentiles.

For simulations where λ_*k*_ = 0 (no change in occupancy resulting from treatment), type 1 error risk was the proportion of times where the credible intervals of 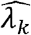 did not overlap 0 (false positive). For all other simulations (there was a change in occupancy resulting from treatment), type 2 error risk was the proportion of times that the credible intervals of 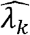 overlapped zero (false negative).

Lastly, we calculated the ‘percent correctly classified’ of the direction of a treatment effect. Managers may not always have the resources to collect sufficiently large datasets that yield precise and accurate species-specific estimates of a treatment effect. They may nevertheless derive value from stating how confident they are that there is *some* effect, and the direction of that effect (helping or harming species). This was calculated as the proportion of the posterior distributions for 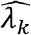 that were correctly classified as positive or negative.

To evaluate sample designs needed for species of concern within a larger community, we recalculated these error metrics using just the species with occupancy and detection intercepts (*β*_0*k*_, *α*_0*k*_)≤ 0.25.

## Results

The multispecies occupancy model (MSOM) was especially sensitive to violations of the species exchangeability assumption (SEA) with smaller datasets. When the SEA was violated (species’ responses were bimodal or randomly distributed), approximately 6 times more data (36 sites, 6-8 surveys) were required to approach similar levels of accuracy for treatment effect estimates as when the SEA was met (same response; 6 sites, 2-3 surveys; Fig. 1). Although the bimodal response may intuitively seem like a stronger violation of the SEA (with two distinct groups), in practice it yielded identical results to the random normal response with respect to estimation error and relative model performance. This is because the average effect magnitudes were equal between these two simulated responses.

**Figure 1.**
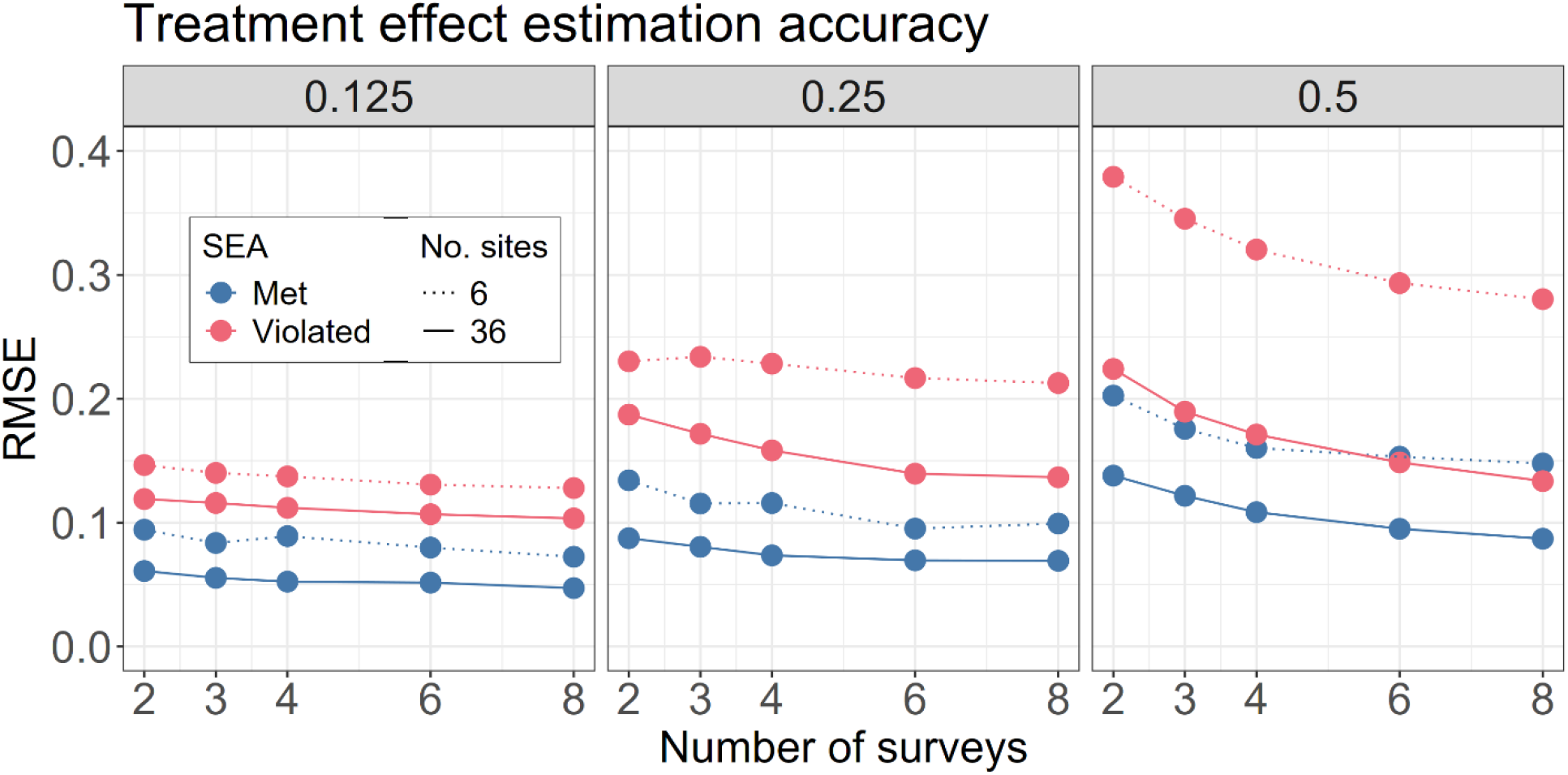
Mean root mean squared error (RMSE) of species-specific treatment effect estimates using the multispecies occupancy model for a community of 50 species when the species exchangeability assumption (SEA) was either met (blue) or violated (red). Three average effect magnitudes shown (columns). Line types represent the number of sites sampled.

In simulated scenarios that met the SEA, the MSOM always outperformed SSOMs and the hybrid model (Fig. 2; refer to supplement Figs. 2a-2f for all community-design combinations). When the SEA was violated, the MSOM yielded less accurate treatment effect estimates than SSOMs or the hybrid model under some low-sampling design configurations. Even so, the differences were relatively small and once 36 sites were sampled, the three models’ error estimates were virtually identical, although RMSE remained higher when SEA was violated than when it was met (Fig. 2, right panel).

**Figure 2.**
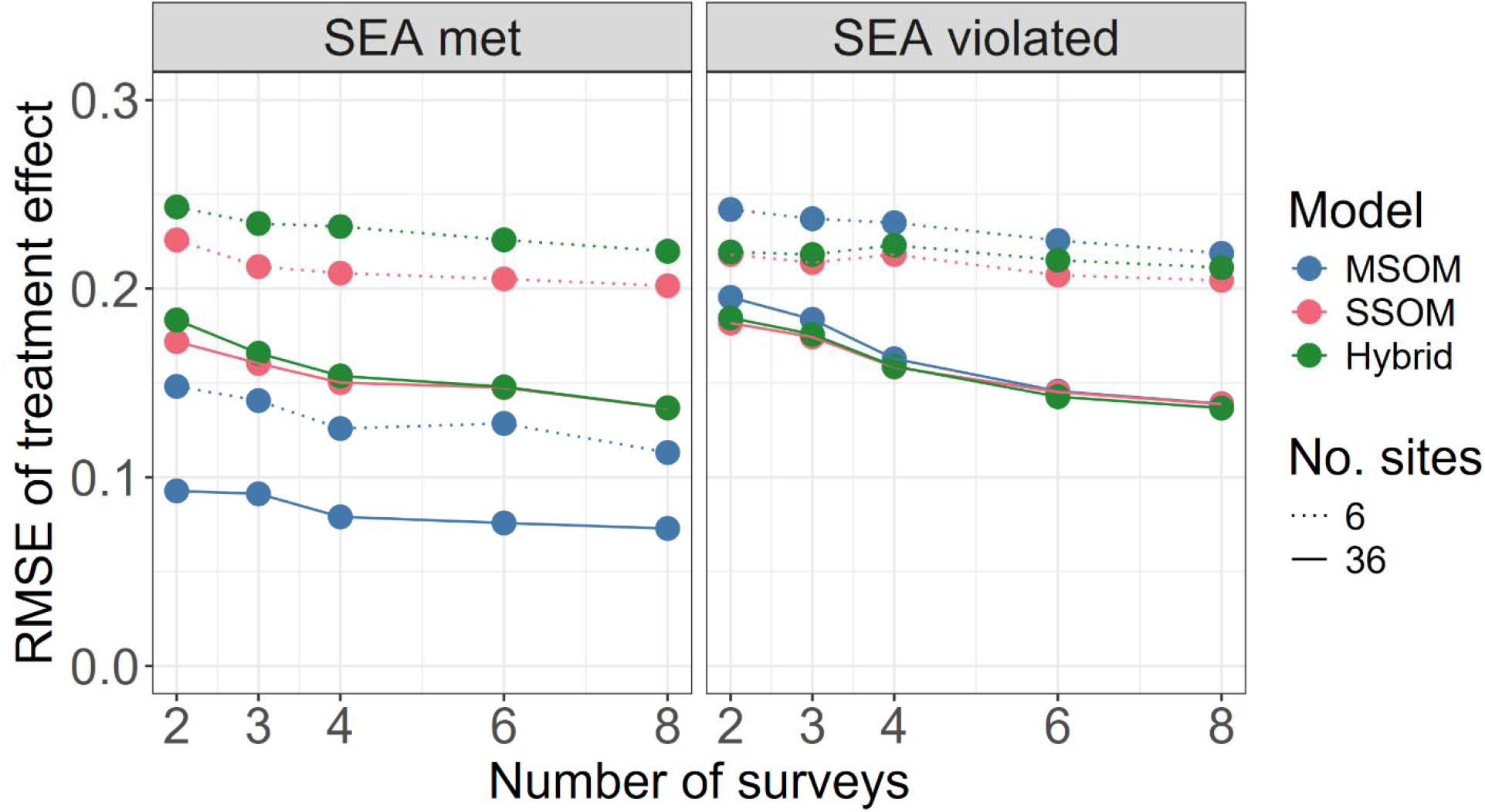
Mean root mean square error (RMSE) for treatment effect estimates when the species exchangeability assumption (SEA) was met (left; all species responded similarly) and violated (right; species responses were bimodal). The multispecies occupancy model (MSOM) minimized error when the SEA was met but otherwise performed similarly to single-species occupancy models (SSOM) and the hybrid model. Shown: communities of 25 species with an intermediate effect magnitude (0.25) sampled across either 6 or 36 sites (line type).

We found little difference in any of the metrics between the two community sizes considered (25 or 50 species; details in supplement). Likewise, recalculating the metrics for species of concern increased error and uncertainty only slightly compared to the entire community (refer to the supplement). Coverage for treatment effect using 95% CIs was generally very good across all models and scenarios considered (refer to supplemental Figs. 3a-3f). However, this was a function of generally large uncertainty (widths) of the CIs (refer to supplemental Figs. 4a-4f) which made this a misleading metric absent additional context. Imprecise treatment effect estimates contributed to low type 1 error risk (inferring some effect when none occurred) and high type 2 error risk (inferring no effect when one occurred). When calculated using the 95^th^ percent CIs, type 1 error risk was always < 4% regardless of model type (refer to supplemental Figs. 5a-5f). Using the 75^th^ percent CI, type 1 error risk under the MSOM never exceeded 20% for the sampling configurations we simulated (refer to supplemental Figs. 6a-6f). Using the 95^th^ percent CI, type 2 error risk tended to range from 0.5 – 1 (Fig. 3). Large average effect magnitudes decreased type 2 error risk, as did meeting the SEA. As a result, using the 75^th^ percent CI may provide more useful information for guiding sampling design (especially considering the relatively low type 1 error risk; refer to supplemental Figs. 8a-8f).

**Figure 3.**
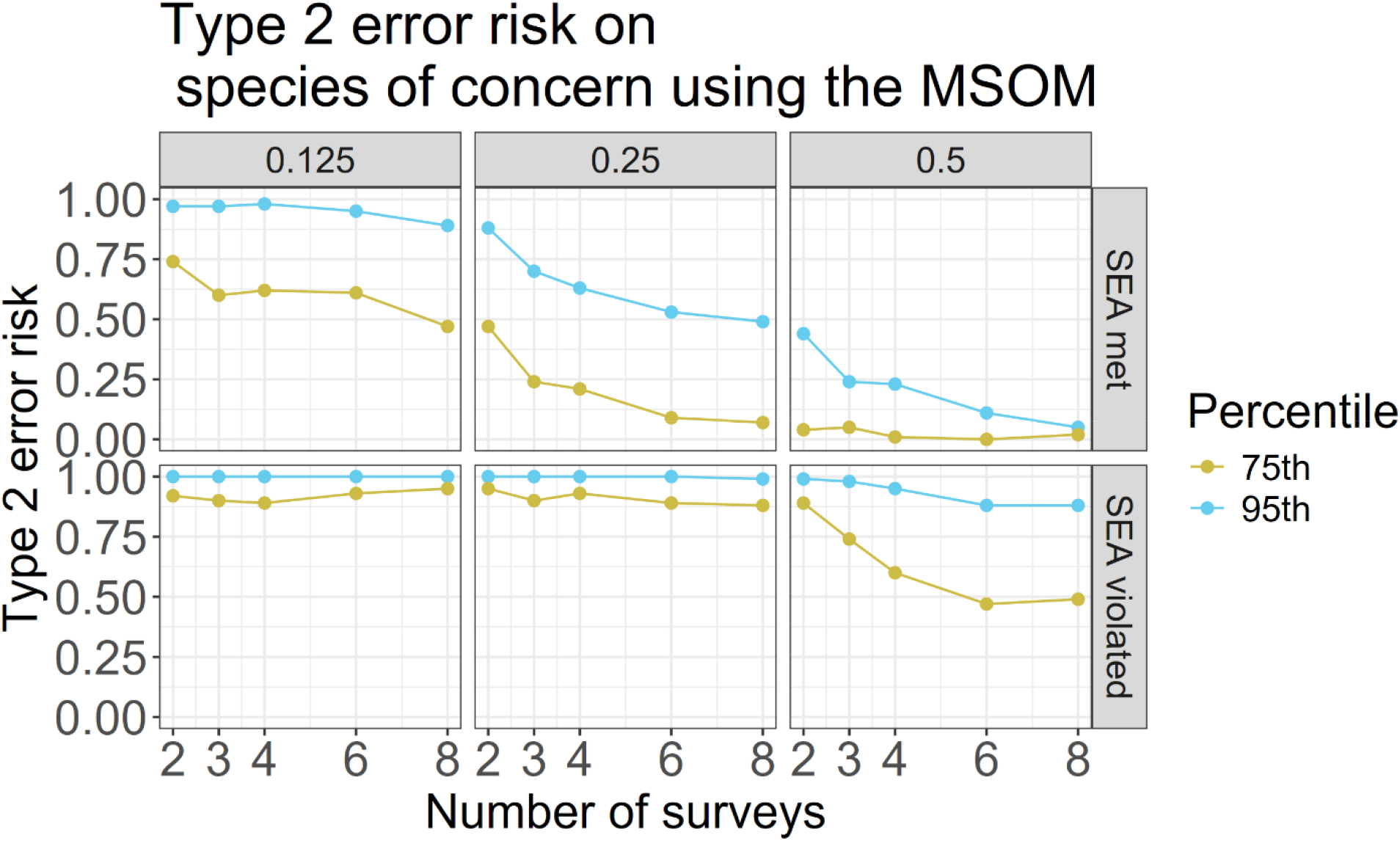
The risk of missing habitat treatment effects on species of concern based on 75^th^ and 95^th^ percent credible intervals overlapping zero (type 2 error) using the multispecies occupancy model (MSOM). Type 2 error risk decreased when the species exchangeability assumption (SEA) was met (top row), effect magnitudes were large (increasing left to right), and more surveys were conducted. Shown: 12 sites sampled, 25 species in the community.

The ‘percent correctly classified’ (PCC) provided an alternative method of quantifying model error based on a binary characterization of treatment (positive or negative effect on occupancy) using the full posterior distributions of species-specific estimates. Despite its simplicity, treatment effects may still be impossible to detect when the SEA is violated and treatment effects are small. Under these conditions, extensive sampling (36 sites, 8 surveys each) can yield marginally higher PCC values using the SSOM (71%) compared to the MSOM (64%; Fig. 4).

**Figure 4.**
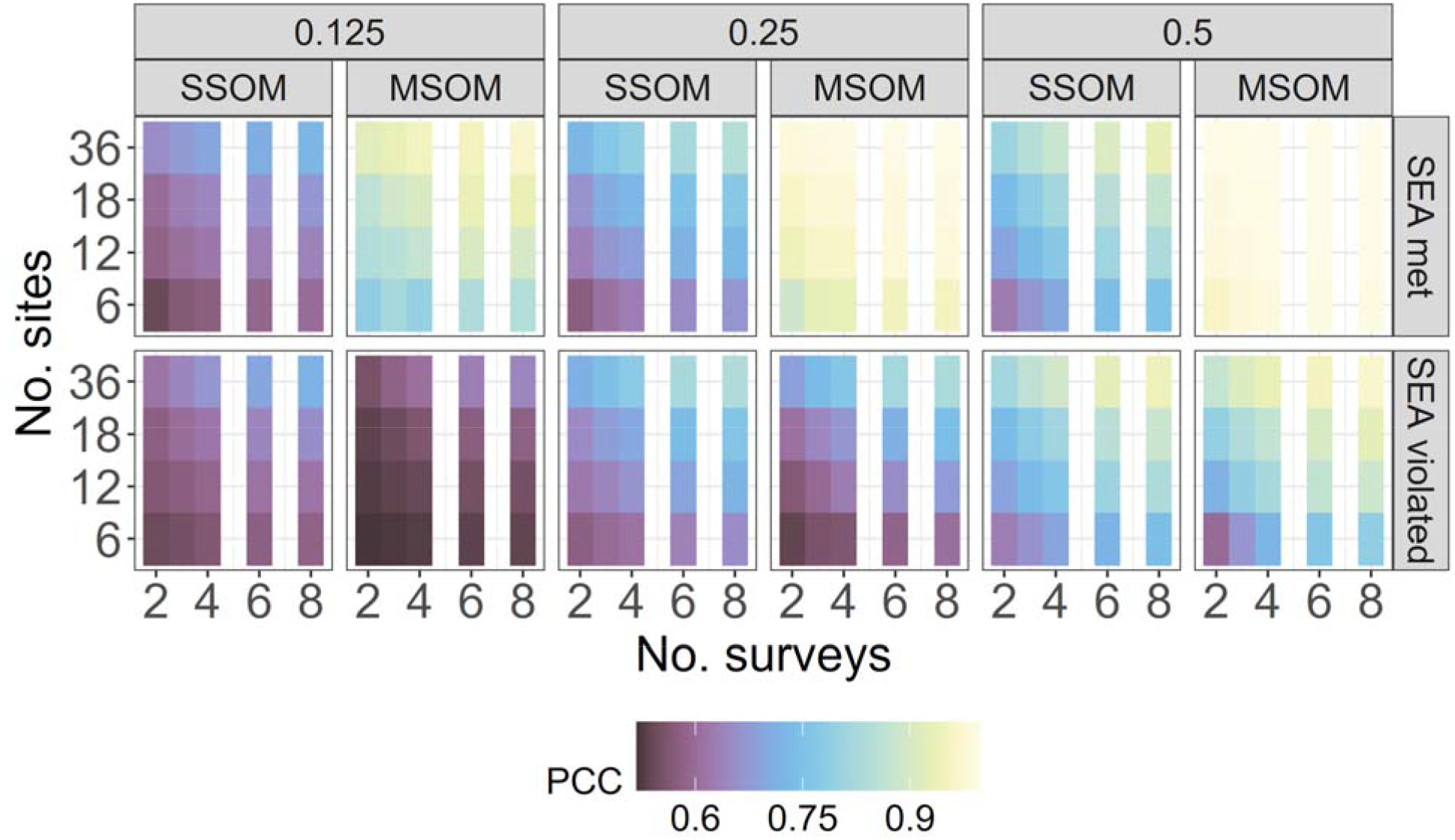
The average percent of species-specific treatment effect posterior distributions that were correctly classified as positive or negative (PCC) using single species occupancy models (SSOM) and the multispecies occupancy model (MSOM). Three effect magnitudes shown (increasing left to right; main columns) when the species exchangeability assumption (SEA) was met (top row) or violated (bottom row). Shown: community size of 50 species.

Tradeoffs between the number of sites and the number of surveys needed to achieve higher PCC values depended primarily on effect magnitude and whether the SEA was met or not (Fig. 4). For instance, when all species respond similarly (SEA met) and with an intermediate effect magnitude, greater than 90% PCC could be achieved with the MSOM through multiple sampling configurations when 12 or more sites were sampled.

## Discussion

When habitat treatments have large effects and influence all species in the community the same way, a moderate amount of sampling effort is required to detect species-specific treatment effects (12 sites and 8 surveys, Fig. 3). In contrast, even moderate habitat treatment effects may be impossible to detect via occupancy modeling using smaller datasets (< 36 sampling locations equally split between treatment/control, < 8 surveys each). The main risk of undersampling was inflated type 2 error – failure to detect meaningful treatment effects on occupancy. The number of species monitored had little impact on treatment response estimation under the conditions we simulated. Although multispecies occupancy models (MSOMs) can drastically improve the accuracy and precision of parameter estimates compared to single species occupancy models (SSOMs), with smaller datasets these improvements rely heavily on the assumption that species respond similarly to treatment (the species exchangeability assumption [SEA]).

Occupancy studies commonly report sampling efforts above the range of conditions we simulated (30 - 100+ sampling locations, 2 - 20+ sampling events; e.g., Zipkin et al. 2012, Broms et al. 2016, Boone et al. 2023). This is somewhat reassuring because larger datasets are more robust to violations of the SEA (Fig 4). However, it is also common to fit models with multiple detection and occupancy covariates. Ideally study design begins with a power analysis tailored to its goals (Devarajan et al., 2020). Yet, public land managers face severe resource constraint limitations (Smiley, 2008). Among their various duties, they are tasked with monitoring rare and threatened species, evaluating the effects of experiment habitat treatments on these species of concern, and making adaptive management decisions (Moore et al., 2011). As such, identifying the minimum amount of data needed to estimate meaningful species-specific effects could support management decisions. Our results and published code can help research support staff in these efforts (Cotterill & Graves, 2025). Study goals – ‘why’ to perform the work and ‘what’ to measure – should be clearly articulated before exploring ‘how’ sampling should proceed (Bailey et al., 2007). Survey design tradeoffs often call for case-by-case consideration and though MSOMs can enhance sampling efficiency, our results show that caution is warranted at lower sampling efforts.

When the goal is quantifying habitat restoration treatment effects on species within a community, ideal species groupings (how the community is defined) would be based on whether species responded positively (increased occupancy) or negatively (decreased occupancy). Yet, this is often what practitioners seek to study, rather than information that is available *a priori*. For pollinator communities – and presumably many other taxa as well – there is little published information with which to justify *a priori* groupings for specific treatments or other environmental covariates. Conceivably raw data could be used to group species’ responses (‘were species observed more times at treatment or control sites?’), but this could be a risky choice if made based on relatively few detections. In other words, for species of concern that are rarely detected, and with fewer surveys and sites, there is substantial risk of assigning those species to the wrong group based on random error. By contrast, when the SEA is met, small treatment effects (+/- 0.125 change in occupancy probability) may be classified (helping or harming) with >90% confidence from as few as 12 sites and 6-8 surveys (Fig. 4).

Detection/non-detection data are, by their nature, simple, and this limits the extent to which model extensions might be expected to improve on the SEA dependence. Therefore, if working with small datasets and in the absence of strong prior information, comparing species-specific estimates using both SSOM and MSOM and performing sensitivity analyses based on grouping decisions with MSOMs may provide insight into violation of the SEA.

In general, the number of species (25 or 50) was far less influential than the range of site and survey conditions we simulated. This is useful information because identifying taxa like insects to species-level often requires specialized training, which is not always readily available. Likewise, eDNA methods cannot always differentiate to the species level. In these cases, species’ detections may be grouped at higher order taxonomic groups (e.g., genus, family) which limits the possible number of taxonomic units (i.e., the number of ‘species’) in the community. Although it forces an additional assumption of exchangeability (that all species within the taxonomic unit are perfectly alike), this may be a necessary approach for some studies – in science and in conservation we often operate from a position of ignorance until we have better information.

Failure to take the species exchangeability assumption into account may lead those in the design stage of monitoring programs to overestimate their statistical power to answer specific questions. When violation of the SEA is expected, sampling at more sites or increasing the number of surveys may be the only way to accurately quantify treatment effects. Multiple options exist in choices to increase sites or surveys based on logistics, though consideration of independence of sites and site similarity should also be considered (Fig. 4). For adaptive management scenarios where some early indication of habitat restoration effects is needed, considering the percent of posterior distributions classified as positive or negative (PCC) may help guide study design, which requires less precision than other commonly used approaches. Other best practices could be to include reporting the full posterior distributions of species-specific parameter values and comparing SSOM and MSOM results.

It is inherently difficult to conduct broadly applicable power analyses because variation across many dimensions generates an intractable number of conditions that must also be performed in replicate. Although we focused on pollinators in the western United States to inform regional goals, the conditions we selected can inform sample sizes needed across many different species groups (e.g., birds, fish) and data collection methods (e.g., eDNA, netting, timed observations). These results provide a strong starting point for study systems where the goal is to efficiently evaluate treatment effects on species of concern. The relative costs associated with any specific project can vary depending on sampling method, the accessibility of sites, or the availability of trained personnel. When specific costs are known, these factors can be formally incorporated into power analyses using optimization procedures (for an example, refer to Sanderlin et al. 2014). The code that supports this analysis has been published to help quantitative support staff evaluate the outcomes for treatments and to help others tailor designs to their needs, for example, investigating greater complexity in the models for occupancy and detection under specific sets of temporal and funding constraints (Cotterill & Graves, 2025).

## Supporting information

Electronic Supplement

## Disclaimer

Any use of trade, firm, or product names is for descriptive purposes only and does not imply endorsement by the U.S. Government.

