## Supplementary material for "Failure to meet the exchangeability assumption in Bayesian multispecies occupancy models: Implications for study design": Electronic Supplement: IP-176524-supplement.html

#### 2025-04-04

- Authors
- Prior
  specification
  - Single species occupancy
    models (SSOMs)
  - Multispecies occupancy
    models (MSOMs)
  - Hybrid model
- Effect types
- Results
  - Root-mean squared error (RMSE)
    - All
      species - SSOM
    - All
      species - MSOM
    - All
      species - Hybrid
    - Only rare species - SSOM
    - Only rare species - MSOM
    - Only rare species - Hybrid
  - Coverage
    - All
      species - SSOM
    - All
      species - MSOM
    - All species - Hybrid
    - Only rare species - SSOM
    - Only rare species - MSOM
    - Only rare species -
      Hybrid
  - CI widths
    - All
      species - SSOM
    - All
      species - MSOM
    - All species - Hybrid
    - Only rare species - SSOM
    - Only rare species - MSOM
    - Only rare species -
      Hybrid
  - Type 1 error risk at 95%
    CI threshold
    - All
      species - SSOM
    - All
      species - MSOM
    - All species - Hybrid
    - Only rare species - SSOM
    - Only rare species - MSOM
    - Only rare species -
      Hybrid
  - Type 1 error risk at 75%
    CI threshold
    - All
      species - SSOM
    - All
      species - MSOM
    - All species - Hybrid
    - Only rare species - SSOM
    - Only rare species - MSOM
    - Only rare species -
      Hybrid
  - Type 2 error risk at 95%
    CI threshold
    - All
      species - SSOM
    - All
      species - MSOM
    - All species - Hybrid
    - Only rare species - SSOM
    - Only rare species - MSOM
    - Only rare species -
      Hybrid
  - Type 2 error risk at 75%
    CI threshold
    - All
      species - SSOM
    - All
      species - MSOM
    - All species - Hybrid
    - Only rare species - SSOM
    - Only rare species - MSOM
    - Only rare species -
      Hybrid
  - Percent correctly classified
    - All
      species - SSOM
    - All
      species - MSOM
    - All species - Hybrid
    - Only rare species - SSOM
    - Only rare species - MSOM
    - Only rare species -
      Hybrid

This information product has been peer reviewed and approved for
publication by the U.S. Geological Survey.

#### Authors

Gavin G. Cotterill^1^, Douglas A. Keinath^2^, and
Tabitha A. Graves^1^

^1^ U.S. Geological Survey, Northern Rocky Mountain Science
Center, 38 Mather Drive, PO Box 169, West Glacier, MT 59936, USA

^2^ U.S. Fish and Wildlife Service, Wyoming Ecological
Services Field Office, 334 Parsley Blvd, Cheyenne, WY 82007, USA

### Prior specification

#### Single species occupancy models (SSOMs)

The species-specific priors were as follows:

\[
\beta\_{0k} \sim dnorm(0, 2.25^{-2}) \\
\beta\_{1k} \sim dnorm(0, 2.25^{-2}) \\
\alpha\_{0k} \sim dnorm(0, 2.25^{-2})
\]

#### Multispecies occupancy models (MSOMs)

The community level hyperpriors were as follows:

\[
\mu\_{\beta0} \sim dnorm(0, 2.25^{-2}) \\
\sigma\_{\beta0} \sim halfCauchy(2) \\
\tau\_{\beta0} = \frac{1}{\sigma\_{\beta0}^2} \\
\mu\_{\beta1} \sim dnorm(0, 2.25^{-2}) \\
\sigma\_{\beta1} \sim halfCauchy(2) \\
\tau\_{\beta1} = \frac{1}{\sigma\_{\beta1}^2} \\
\mu\_{\alpha0} \sim dnorm(0, 2.25^{-2}) \\
\sigma\_{\alpha0} \sim halfCauchy(2) \\
\tau\_{\alpha0} = \frac{1}{\sigma\_{\alpha0}^2}
\]

The species-specific priors were as follows:

\[
\beta\_{0k} \sim dnorm(\mu\_{\beta0}, \tau\_{\beta0}) \\
\beta\_{1k} \sim dnorm(\mu\_{\beta1}, \tau\_{\beta1}) \\
\alpha\_{0k} \sim dnorm(\mu\_{\alpha0}, \tau\_{\alpha0})
\]

#### Hybrid model

The community-level hyperpriors were as follows:

\[
\mu\_{\beta0} \sim dnorm(0, 2.25^{-2}) \\
\sigma\_{\beta0} \sim halfCauchy(2) \\
\tau\_{\beta0} = \frac{1}{\sigma\_{\beta0}^2} \\
\mu\_{\alpha0} \sim dnorm(0, 2.25^{-2}) \\
\sigma\_{\alpha0} \sim halfCauchy(2) \\
\tau\_{\alpha0} = \frac{1}{\sigma\_{\alpha0}^2}\\
\]

The species-level priors were as follows:

\[
\beta\_{0k} \sim dnorm(\mu\_{\beta0}, \tau\_{\beta0}) \\
\beta\_{1k} \sim dnorm(0, 2.25^{-2}) \\
\alpha\_{0k} \sim dnorm(\mu\_{\alpha0}, \tau\_{\alpha0})
\]

### Effect types

Figure 1. Histograms of species-specific treatment effects on the
probability scale at three levels of magnitude (columns) under three
scenarios (rows). Under the single and two-guild scenarios treatment
effect increased or decreased occupancy by 0.125, 0.25, or 0.5.
Occupancy cannot exceed 1. Species with high occupancy intercepts
(>|.875|, >|.75|, >|.5|) were assigned treatment effects that
resulted in saturation at treatment sites (summed to one). As a result,
mean effect magnitudes were less than the targeted increase, and were
approximately 0.12, 0.22, and 0.37 for the three effect types.

### Results

#### Root-mean squared error (RMSE)

\[
rmse = \sqrt\frac{\sum(\hat\lambda\_k - \lambda\_k)^2}{100 \* nspp}
\] Where *nspp* was the community size and there were 100
replicates for each study combination.

##### All species - SSOM

Figure 2a. Comparing root mean squared error of estimated treatment
effect for 100 replicates at each design X simulation combination when
single-species occupancy models were fit for all species in the
community.

##### All species - MSOM

Figure 2b. Comparing root mean squared error of estimated treatment
effect for 100 replicates at each design X simulation combination under
the multispecies occupancy model.

##### All species - Hybrid

Figure 2c. Comparing root mean squared error of estimated treatment
effect for 100 replicates at each design X simulation combination under
the hybrid model.

##### Only rare species - SSOM

Figure 2d. Comparing root mean squared error of estimated treatment
effect for 100 replicates at each design X simulation combination when
single-species occupancy models were fit for all species in the
community (rare species only).

##### Only rare species - MSOM

Figure 2e. Comparing root mean squared error of estimated treatment
effect for 100 replicates at each design X simulation combination under
the multispecies occupancy model (rare species only).

##### Only rare species - Hybrid

Figure 2f. Comparing root mean squared error of estimated treatment
effect for 100 replicates at each design X simulation combination under
the hybrid model (rare species only).

#### Coverage

Coverage was defined as the proportion of the time that the true
treatment effect (\(\hat\lambda\_k\))
fell within the 95% CI of treatment effect estimates (\(\lambda\_k\)).

##### All species - SSOM

Figure 3a. 95% confidence interval coverage of treatment effect
estimates under the single species occupancy model (SSOM).

##### All species - MSOM

Figure 3b. 95%
confidence interval coverage of treatment effect estimates under the
multispecies occupancy model (MSOM).

##### All species - Hybrid

Figure 3c. 95%
confidence interval coverage of treatment effect estimates under the
hybrid occupancy model.

##### Only rare species - SSOM

Figure 3d. 95% confidence interval coverage of treatment effect
estimates under the single species occupancy model (SSOM) when only
species of concern were considered.

##### Only rare species - MSOM

Figure 3e. 95% confidence interval coverage of treatment effect
estimates under the multispecies occupancy model (MSOM) when only
species of concern were considered.

##### Only rare species - Hybrid

Figure 3f. 95% confidence interval coverage of treatment effect
estimates under the hybrid occupancy model when only species of concern
were considered.

#### CI widths

Coverage values were high because the confidence intervals were very
wide. The widths mostly change as a function of the number of surveys,
and change less as a function of effect magnitude.

##### All species - SSOM

Figure 4a. 95% confidence interval widths (probability scale) for
treatment effect estimates under the single species occupancy model
(SSOM).

##### All species - MSOM

Figure 4b. 95% confidence interval widths (probability scale) for
treatment effect estimates under the multispecies occupancy model
(MSOM).

##### All species - Hybrid

Figure 4c. 95% confidence interval widths (probability scale) for
treatment effect estimates under the hybrid occupancy model.

##### Only rare species - SSOM

Figure 4d. 95% confidence interval widths (probability scale) for
treatment effect estimates under the single species occupancy model
(SSOM) when only species of concern were considered.

##### Only rare species - MSOM

Figure 4e. 95% confidence interval widths (probability scale) for
treatment effect estimates under the multispecies occupancy model (MSOM)
when only species of concern were considered.

##### Only rare species - Hybrid

Figure 4f. 95% confidence interval widths (probability scale) for
treatment effect estimates under the hybrid occupancy model when only
species of concern were considered.

#### Type 1 error risk at 95% CI threshold

The proportion of the time that the 95% CI of treatment effect does
not overlap zero when there is no effect in the simulated data, but we
nevertheless fit models that estimate an effect.

##### All species - SSOM

Figure 5a. Type 1 error risk under the single species occupancy model
(SSOM).

##### All species - MSOM

Figure 5b. Type 1 error risk under the multispecies occupancy model
(MSOM).

##### All species - Hybrid

Figure 5c. Type 1 error risk under the hybrid occupancy model.

##### Only rare species - SSOM

Figure 5d. Type 1 error risk under the single species occupancy model
(SSOM) when only species of concern were considered.

##### Only rare species - MSOM

Figure 5e. Type 1 error risk under the multispecies occupancy model
(MSOM) when only species of concern were considered.

##### Only rare species - Hybrid

Figure 5f. Type 1 error risk under the hybrid occupancy model when
only species of concern were considered.

#### Type 1 error risk at 75% CI threshold

The proportion of the time that the 75% CI of treatment effect does
not overlap zero when there is no effect in the simulated data, but we
nevertheless fit models that estimate an effect.

##### All species - SSOM

Figure 6a. Type 1 error risk under the single species occupancy model
(SSOM).

##### All species - MSOM

Figure 6b. Type 1 error risk under the multispecies occupancy model
(MSOM).

##### All species - Hybrid

Figure 6c. Type 1 error risk under the hybrid occupancy model.

##### Only rare species - SSOM

Figure 6d. Type 1 error risk under the single species occupancy model
(SSOM) when only species of concern were considered.

##### Only rare species - MSOM

Figure 6e. Type 1 error risk under the multispecies occupancy model
(MSOM) when only species of concern were considered.

##### Only rare species - Hybrid

Figure 6f. Type 1 error risk under the hybrid occupancy model when
only species of concern were considered.

#### Type 2 error risk at 95% CI threshold

The proportion of the time that the 95% CI of treatment effect
overlaps zero when there is a true effect in the simulated data.

##### All species - SSOM

Figure 7a. Type 2 error risk under the single species occupancy model
(SSOM).

##### All species - MSOM

Figure 7b. Type 2 error risk under the multispecies occupancy model
(MSOM).

##### All species - Hybrid

Figure 7c. Type 2 error risk under the hybrid occupancy model.

##### Only rare species - SSOM

Figure 7d. Type 2 error risk under the single species occupancy model
(SSOM) when only species of concern were considered.

##### Only rare species - MSOM

Figure 7e. Type 2 error risk under the multispecies occupancy model
(MSOM) when only species of concern were considered.

##### Only rare species - Hybrid

Figure 7f. Type 2 error risk under the hybrid occupancy model when
only species of concern were considered.

#### Type 2 error risk at 75% CI threshold

The proportion of the time that the 75% CI of treatment effect
overlaps zero when there is a true effect in the simulated data.

##### All species - SSOM

Figure 8a. Type 2 error risk under the single species occupancy model
(SSOM).

##### All species - MSOM

Figure 8b. Type 2 error risk under the multispecies occupancy model
(MSOM).

##### All species - Hybrid

Figure 8c. Type 2 error risk under the hybrid occupancy model.

##### Only rare species - SSOM

Figure 8d. Type 2 error risk under the single species occupancy model
(SSOM) when only species of concern were considered.

##### Only rare species - MSOM

Figure 8e. Type 2 error risk under the multispecies occupancy model
(MSOM) when only species of concern were considered.

##### Only rare species - Hybrid

Figure 8f. Type 2 error risk under the hybrid occupancy model when
only species of concern were considered.

#### Percent correctly classified

For each species-specific treatment effect estimate in each model
run, the proportions of the posterior distributions falling on the
‘correct side’ of zero were averaged across the community and 100
iterations.

##### All species - SSOM

Figure 9a. Percent of correctly classified treatment effect posterior
distributions under the single species occupancy model (SSOM).

##### All species - MSOM

Figure 9b. Percent of correctly classified treatment effect posterior
distributions under the multispecies occupancy model (MSOM).

##### All species - Hybrid

Figure 9c. Percent of correctly classified treatment effect posterior
distributions under the hybrid occupancy model.

##### Only rare species - SSOM

Figure 9d. Percent of correctly classified treatment effect posterior
distributions under the single species occupancy model (SSOM) when only
species of concern were considered.

##### Only rare species - MSOM

Figure 9e. Percent of correctly classified treatment effect posterior
distributions under the multispecies occupancy model (MSOM) when only
species of concern were considered.

##### Only rare species - Hybrid

Figure 9f. Percent of correctly classified treatment effect posterior
distributions under the hybrid occupancy model when only species of
concern were considered.

Disclaimer: Any use of trade, firm, or product names is for
descriptive purposes only and does not imply endorsement by the U.S.
Government.
